# Genomic sequencing of the aquatic *Fusarium* spp. QHM and BWC1 and their potential application in environmental protection

**DOI:** 10.1101/659755

**Authors:** Hongfei Zhu, Long Zhu, Ning Ding

**Author notes:** Address correspondence to Hongfei Zhu,.

## Abstract

*Fusarium* species are distributed widely in ecosystems of a wide pH range and play a pivotal role in the aquatic community through the degradation of xenobiotic compounds and secretion of secondary metabolites. The elucidation of their genome would therefore be highly impactful with regard to the control of environmental pollution. Therefore, in this study, two indigenous strains of aquatic *Fusarium*, QHM and BWC1, were isolated from a coal mine pit and a subterranean river respectively, cultured under acidic conditions, and sequenced. Phylogenetic analysis of these two isolates was conducted based on the sequences of internal transcript (ITS1 and ITS4) and encoding β-microtubulin (TUB2), translation elongation factors (TEFs) and the second large sub-unit of RNA polymerase (RPB2). *Fusarium*, QHM could potentially represent a new species within the *Fusarium fujikuroi* species complex. *Fusarium* BWC1 were found to form a clade with *Fusarium subglutinans* NRRL 22016, and predicted to be *Fusarium subglutinans.* Shot-gun sequencing on the Illumina Hiseq×10 Platform was used to elucidate the draft genomes of the two species. Gene annotation and functional analyses revealed that they had bio-degradation pathways for aromatic compounds; further, their main pathogenic mechanism was found to be the efflux pump. To date, the genomes of only a limited number of acidic species from the *Fusarium fujikuroi* species complex, especially from the aquatic species, have been sequenced. Therefore, the present findings are novel and have important potential for the future in terms of environmental control.

**IMPORTANCE:** Fusarium genus has over 300 species and were distributed in a variety of ecosystem. Increasing attention has been drawn to *Fusarium* due to the importance in aquatic community, pathogenicity and environmental protection. The genomes of the strains in this work isolated in acidic condition, were sequenced. The analysis has indicated that the isolates were able to biodegrade xenobiotics, which makes it potentially function as environmental bio-agent for aromatic pollution control and remediation. Meanwhile, the virulence and pathogenicity were also predicted for reference of infection control. The genome information may lay foundation for the fungal identification, disease prevention resulting from these isolates and other “-omics” research. The isolates were phylogenetically classified into *Fusarium fujikuroi* species complex by means of concatenated gene analysis, serving as new addition to the big complex.

## Introduction

Species of the genus *Fusarium* are an important group of fungi that are distributed widely in below-ground and above-ground habitats (1, 2). Certain *Fusarium* species can even be found in aquatic habitats, including coal mine pits and subterranean rivers (2). These species are also characterized by their ability to grow in environments with a wide pH range (1, 2). The genus *Fusarium* is important from the perspective of environmental protection and phytopathogenicity: On the one hand, *Fusarium* spp. have potential as bio-control agents, on account of their ability to degrade a variety of xenobiotics, such as aromatic compounds from water or soil. On the other hand, they infect a broad spectrum of crops, thereby resulting in huge economic losses or mycotoxin contamination (3). Given the complex characteristics of these species, it is imperative to elucidate the *Fusarium* genome sequence for the elimination of pollutants while ensuring the control of its pathogenicity. Further, as the *Fusarium* genus forms a “species complex” with the *Aspergillus* genus on account of closely related species, species-level identification of *Fusarium* is also necessary (4). Whole-genome sequencing would be ideal, as this method provides more information than other sequencing approaches in terms of the elucidation of acidic mechanisms, biodegradation and microbial identification. Fungus-specific genes, such as *ITS1/ITS4*, *TUB2*, *TEF* and *RPB2*, and their sequences have been universally used for species identification and analysis of evolutionary relationships (4). The simultaneous use of the fungus-specific genes may phylogenetically give more resolution to over 90% gene identity within *Fusarium* species complex.

The *Fusarium* spp. QHM and BWC1 have been isolated and purified from the 9KG medium culture in the process of isolation of *Acidiphilium cryptum* from coal mine water and underground river-water. Based on the isolation environment, *Fusarium* spp. QHM and BWC1 were thought to be indigenous and predicted to the best solution for the *in situ* remediation of acidic coal mine water (5). These fungal isolates were experimentally proved to be tolerant in a wide pH range of 3.0 to 8.5; therefore, they also have the potential to act as bio-agents for the treatment of acid mine drainage (5, 6). Accordingly, the genomes of both *Fusarium* QHM and *Fusarium* BWC1 isolates were sequenced, annotated and analyzed using bioinformatics methods. The information obtained is highly valuable, as genomic sequences from aquatic fungal isolates, especially from coal mine water, are scant. The yielded genomic sequences and functions will prove useful for research endeavors in the field of “-omics”, microbial communities and water pollution management. Importantly, this genomic information could be used to investigate the biodegradation of aromatic compounds, such phenol and benzo[a]pyrene, for water pollution control and bio-remediation of soil contaminated with polycyclic aromatic hydrocarbons (7).

In this work, we present two draft genomes of *Fusarium* spp. QHM and BWC1. Based on the genes sequenced and their annotational and functional analysis, the evolutionary relationships between these species and their aromatic metabolism were investigated. The findings are expected to have future applications in the management and remediation of coal mine water or underground river water.

## Results

### 1. Isolation of *Fusarium* spp. from the water samples

Two isolates, *Fusarium* spp. QHM and BWC1, were obtained from the water samples. These strains exhibit growth that is visible to the naked eye when cultured in 9 KG medium at a pH of 3.5 to 8.5. The optimum pH for growth is 5.0 to 6.0, so these species are considered to be acidophiles. The overall pH tendancy during the growth was to slide down to lower level in the first two or three days. From inoculation to the exponential stage, pH 5.0-6.0 may go down to pH 2.5-3.5. *Fusarium* sp. QHM and *Fusarium* sp. BWC1 are currently stored in the in-house laboratory of the university.

### 2. Characterization

The sequences obtained after PCR amplification of the ITS1, TUB2, TEF and RPB2 genes were submitted to the GenBank of the National Centre of Biological Information (NCBI) under accession No. MK791252, MK850849, MK850848, MK850847, MK898823, MK907693, MK907692 and MK907691. The BLAST results are provided in Table 1.

**Table 1.**
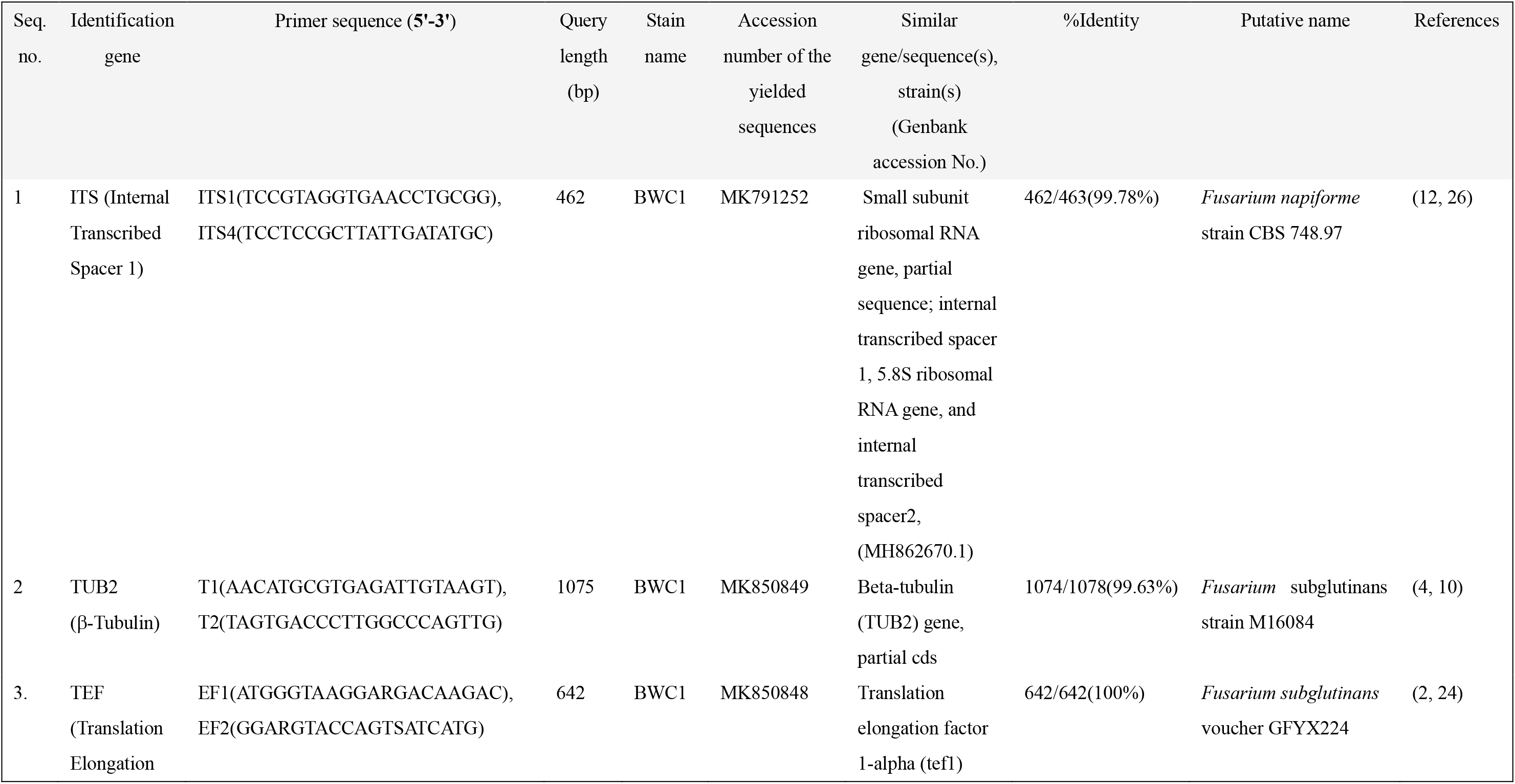
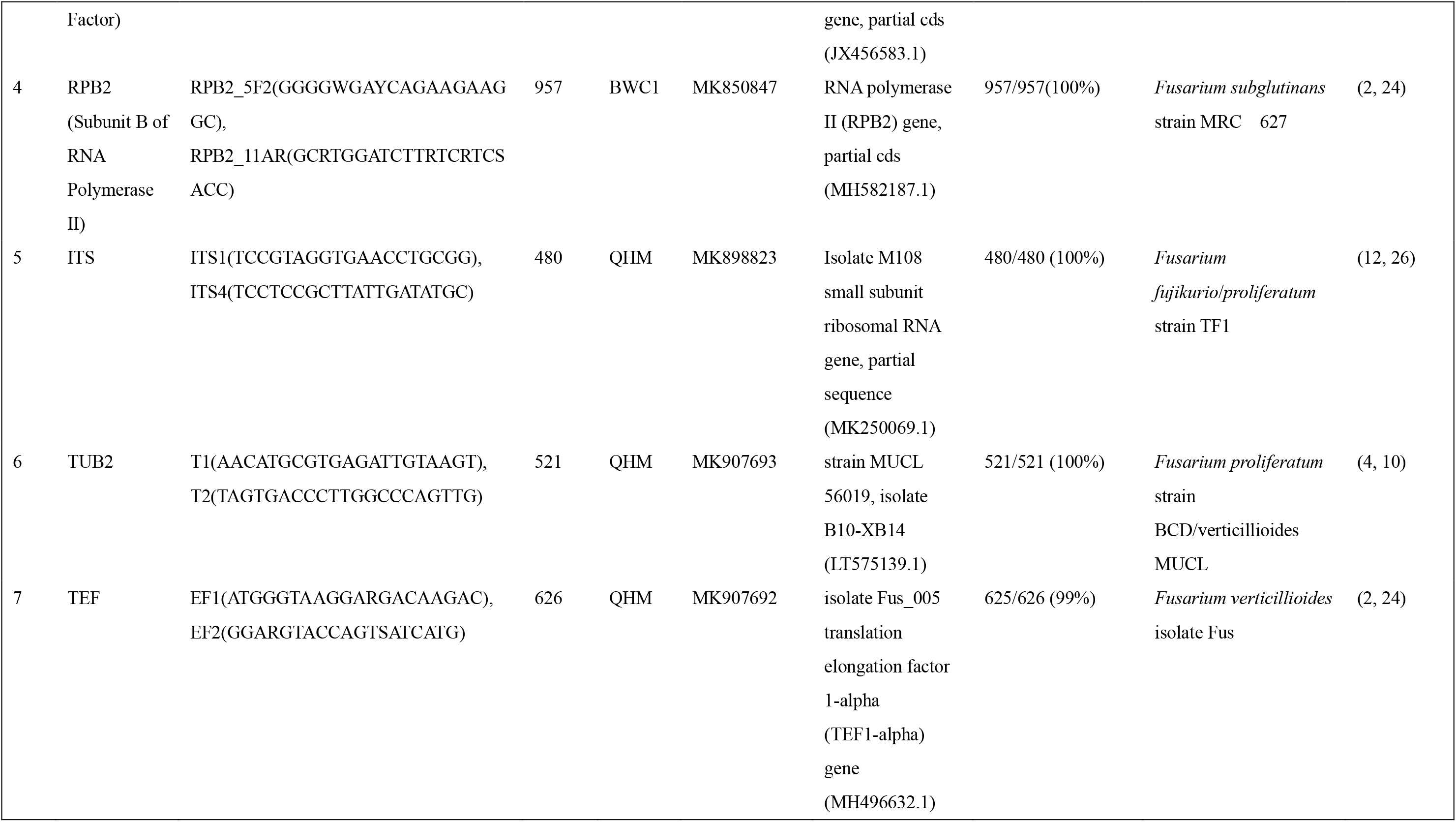
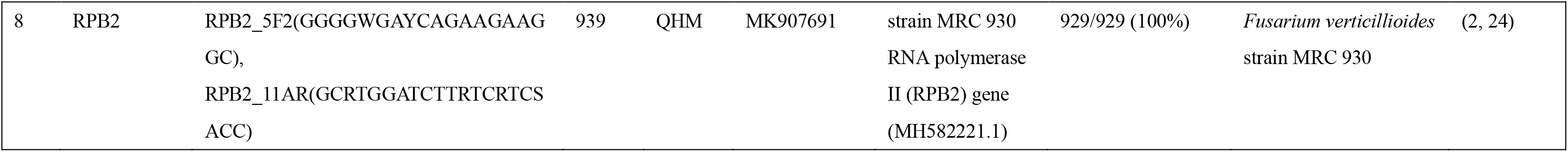
Identification genes and characteristics of the yielded sequences of the analyzed genes in *Fusarium* spp. QHM and BWC1

### 3. Sequencing data

The Whole Genome Shotgun project sequences have been deposited in GenBank under accession numbers SWCP00000000 (*Fusarium* sp. QHM) and SWCQ00000000 (*Fusarium* sp. BWC1). In total, 479 and 2352 scaffolds were obtained for *Fusarium* spp. QHM and BWC1 respectively. The overall results of the assemblage are summarized in Table 2. The GC content of the genomic sequences of both species was 44% to 60%, which is in line with the sequencing data. The predicted coding genes are shown in Table 3.

**Table 2.**
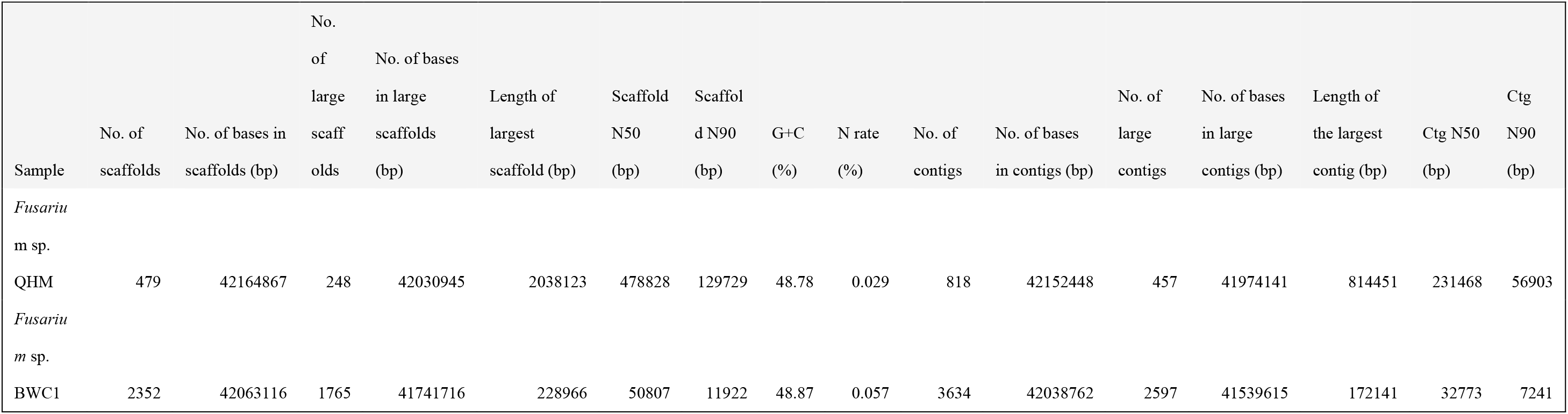
Overall results of genome assemblage

**Table 3.**
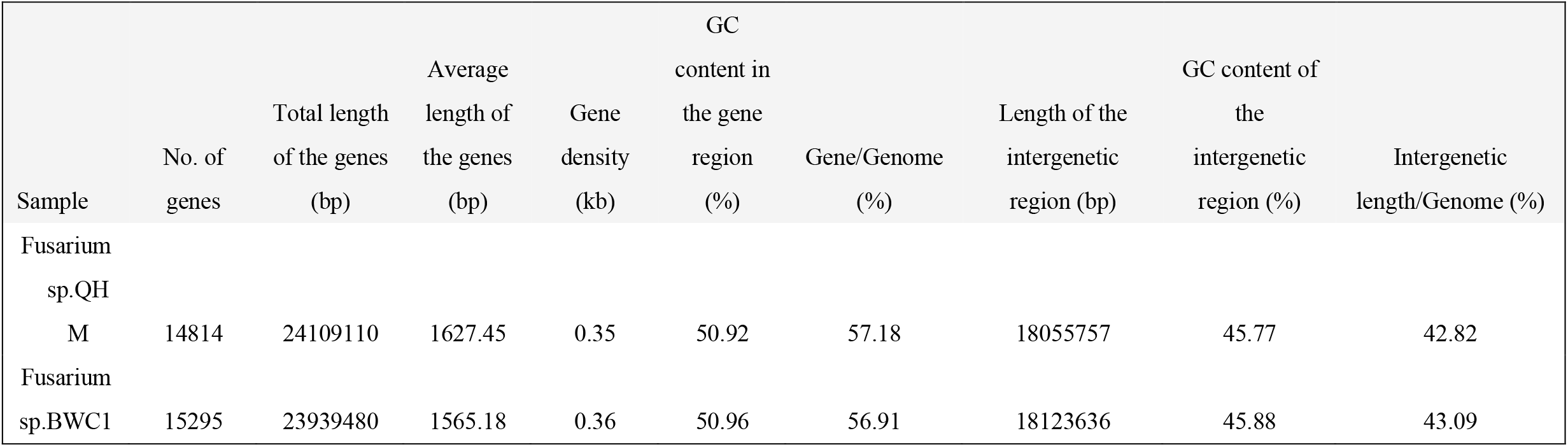
Predictive functions of the coding genes of *Fusarium* spp. QHM and BWC1

### 4. Gene annotation results

The gene annotation results for *Fusarium* spp. QHM and BWC1 were similar, which suggested that they may evolve from a later common ancestor because of a speciation. The minor divergence derived from duplication. As shown in Figure 1, based on Gene Ontology (GO) analysis, both genes were categorized into three main groups that were further divided into various subcategories: biological processes, cellular components and molecular functions. The majority of genes were assigned to metabolic processes, and the second highest number of genes were assigned to catalytic activity. The findings indicate that genes involved in primary and secondary metabolic processes, including secretion, degradation and catalysis, were identified in both strains. Information from the five databanks was used as reference to determine the functions of all the genes from both samples. The annotation results are summarized in Figure 1. The fundamental annotations for both strains in the five public databanks are provided in two excel files, Table S1 and Table S2 in Supplemental Material.

**Figure 1.**
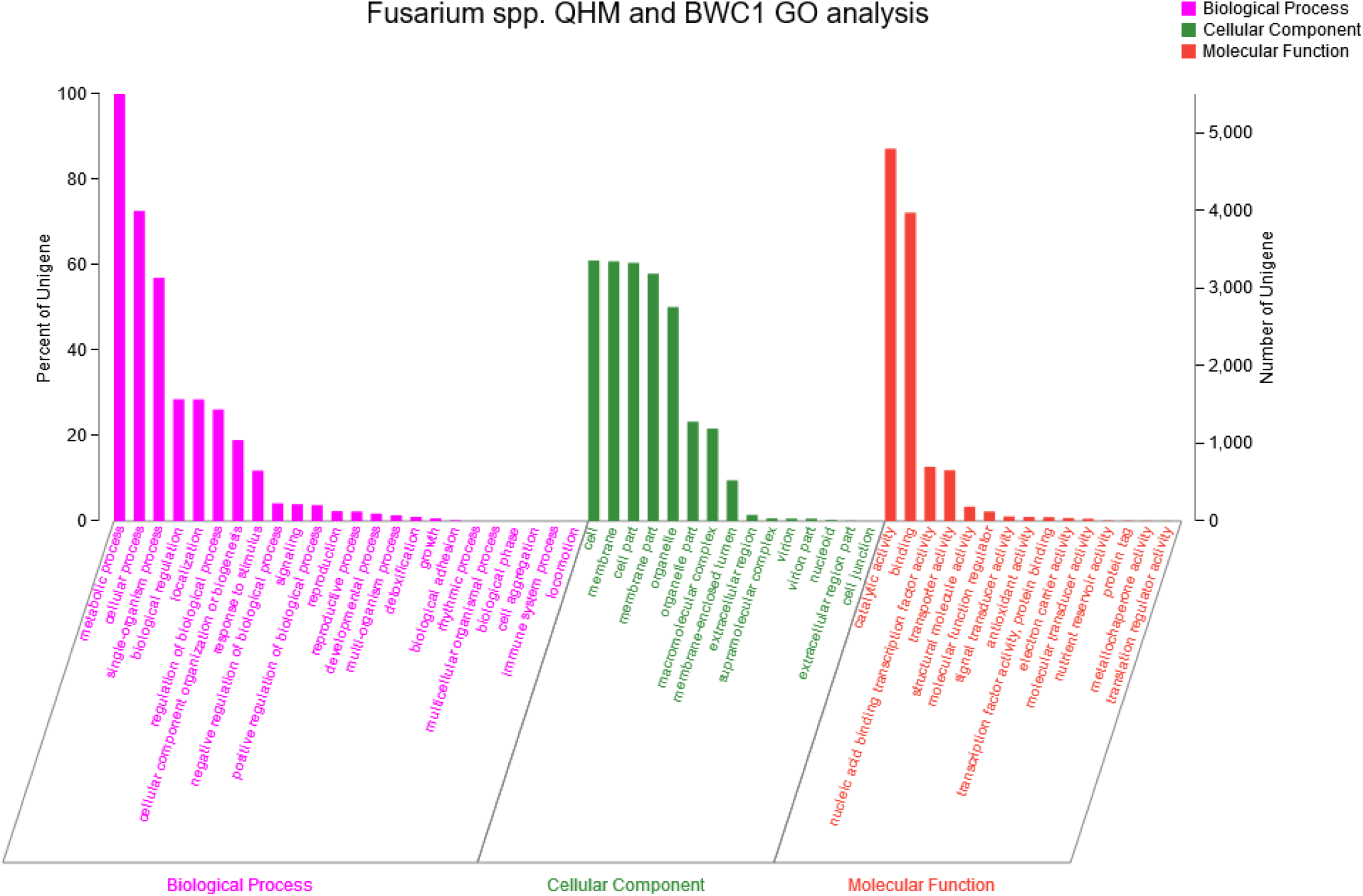
*Fusarium* spp. QHM and BWC1 bar charts obtained from GO analysis. The gene orthologs of *Fusarium* spp. QHM and BWC1 are similar generally. All genes were categorized into three main groups that were further divided into various subcategories: biological processes, cellular components and molecular functions. The majority of genes belongs to metabolic processes, and the second highest number of genes were assigned to catalytic activity group as usual.

Histograms of the findings obtained from Kyoto Encyclopedia of Genes and Genomes (KEGG) pathway analysis, as shown in Figure 2 and 3, show the classification of the genes from both species into six categories: metabolism, human disease, organismal systems, genetic information processing, environmental information processing and cellular processes. The number of genes involved in metabolic processes was 3243 and 3423 in *Fusarium* spp. QHM and BWC1 respectively. The main pathways identified were xenobiotic biodegradation, carbohydrate metabolism and amino acid metabolism. Gene enrichment analysis of the transport- and catabolism-related genes indicated that a high number of genes were involved in these functions at the cellular level. Further, the number of translation-associated genes identified in both species indicate that these genes played a dynamic role in genetic information processing in both strains.

**Figure 2.**
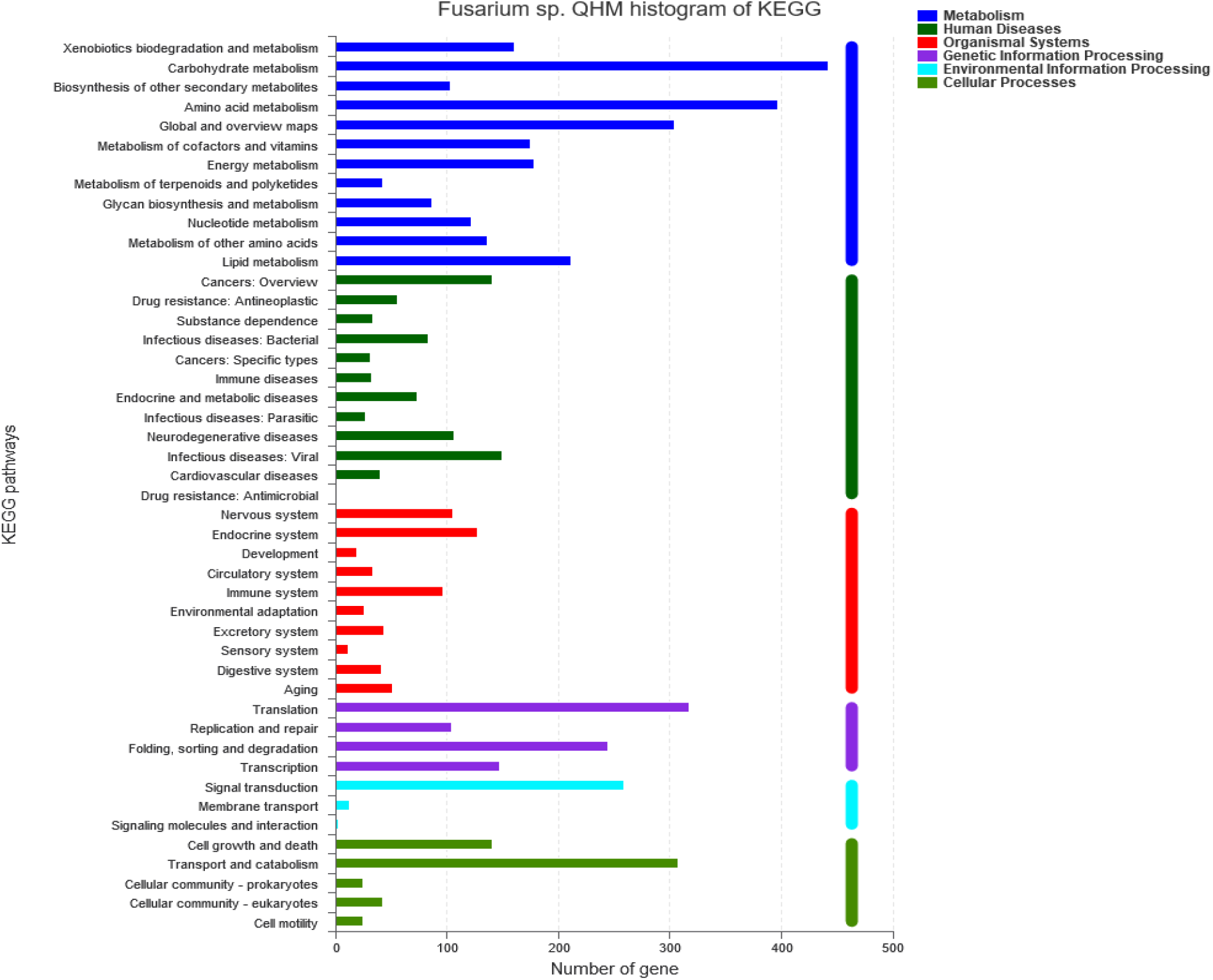
*Fusarium* sp. QHM histogram obtained from KEGG data.

**Figure 3.**
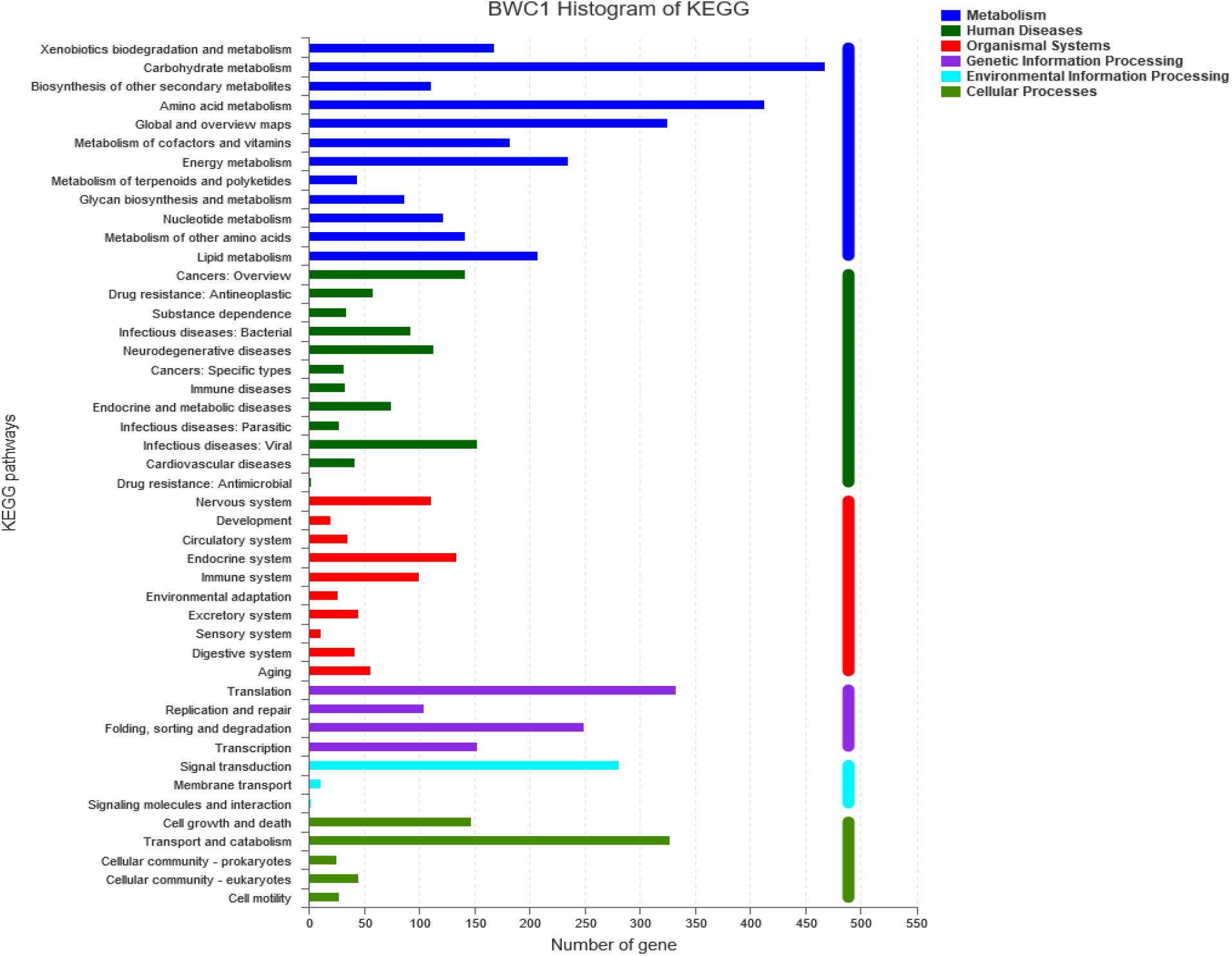
*Fusarium* sp. BWC1 histogram obtained from KEGG data.

There are six main enzyme families that are involved in the synthesis, metabolism and recognition of complex carbohydrates: glycoside hydrolases, glycosyltransferases, polysaccharide lyases, carbohydrate esterases and non-catalytic carbohydrate-binding modules. Carbohydrate hydrolases, which form the majority of the enzyme components, comprise 41.87% and 41.88% of the genes associated with complex carbohydrates in *Fusarium* spp. QHM and BWC1, respectively, as shown in Figure 4 and 5. Overall, the difference between *Fusarium* spp. QHM and BWC1 genomes with regard to these six gene families is inconspicuous.

**Figure 4.**
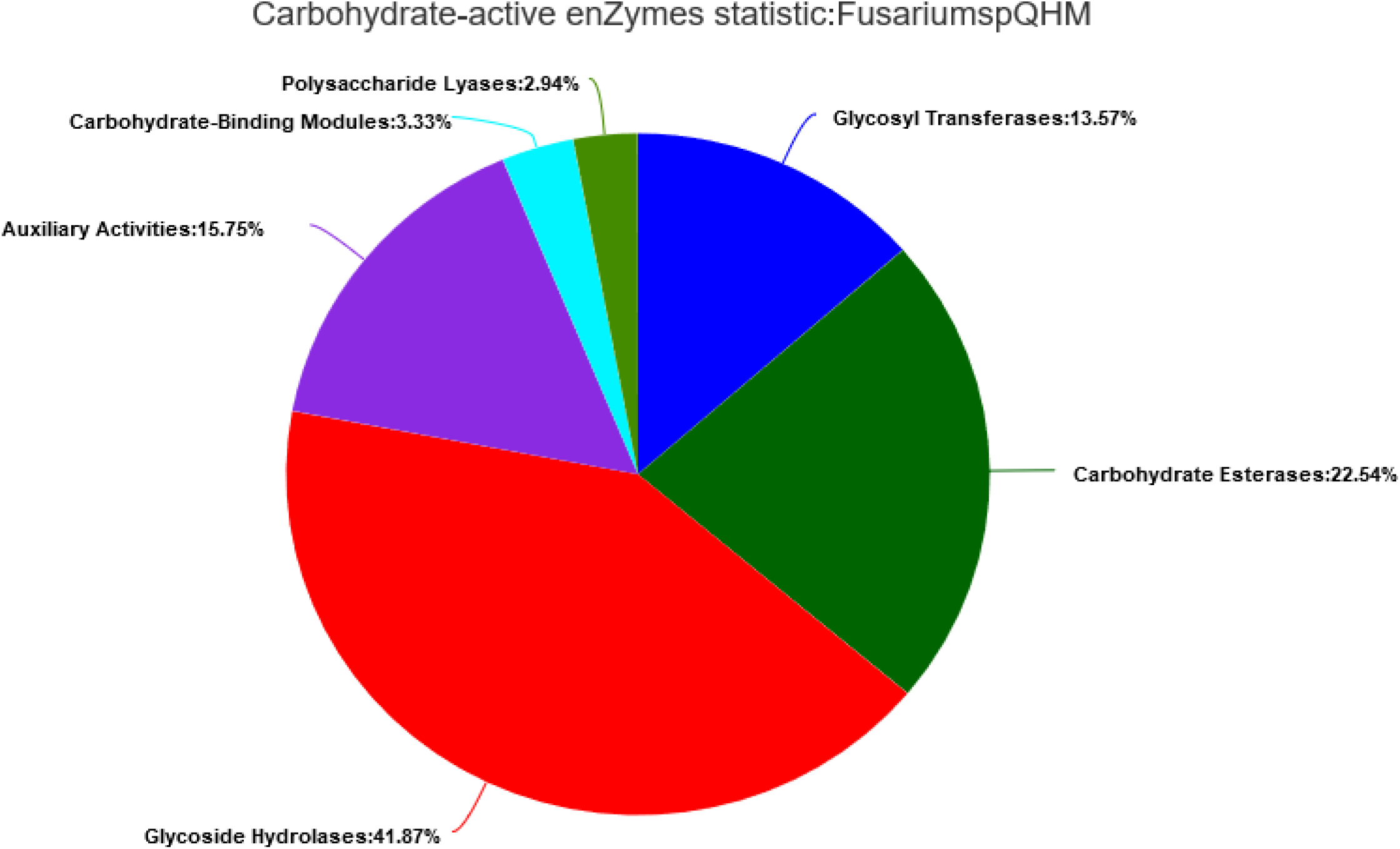
Fusarium sp. QHM CAZy analysis chart.

**Figure 5.**
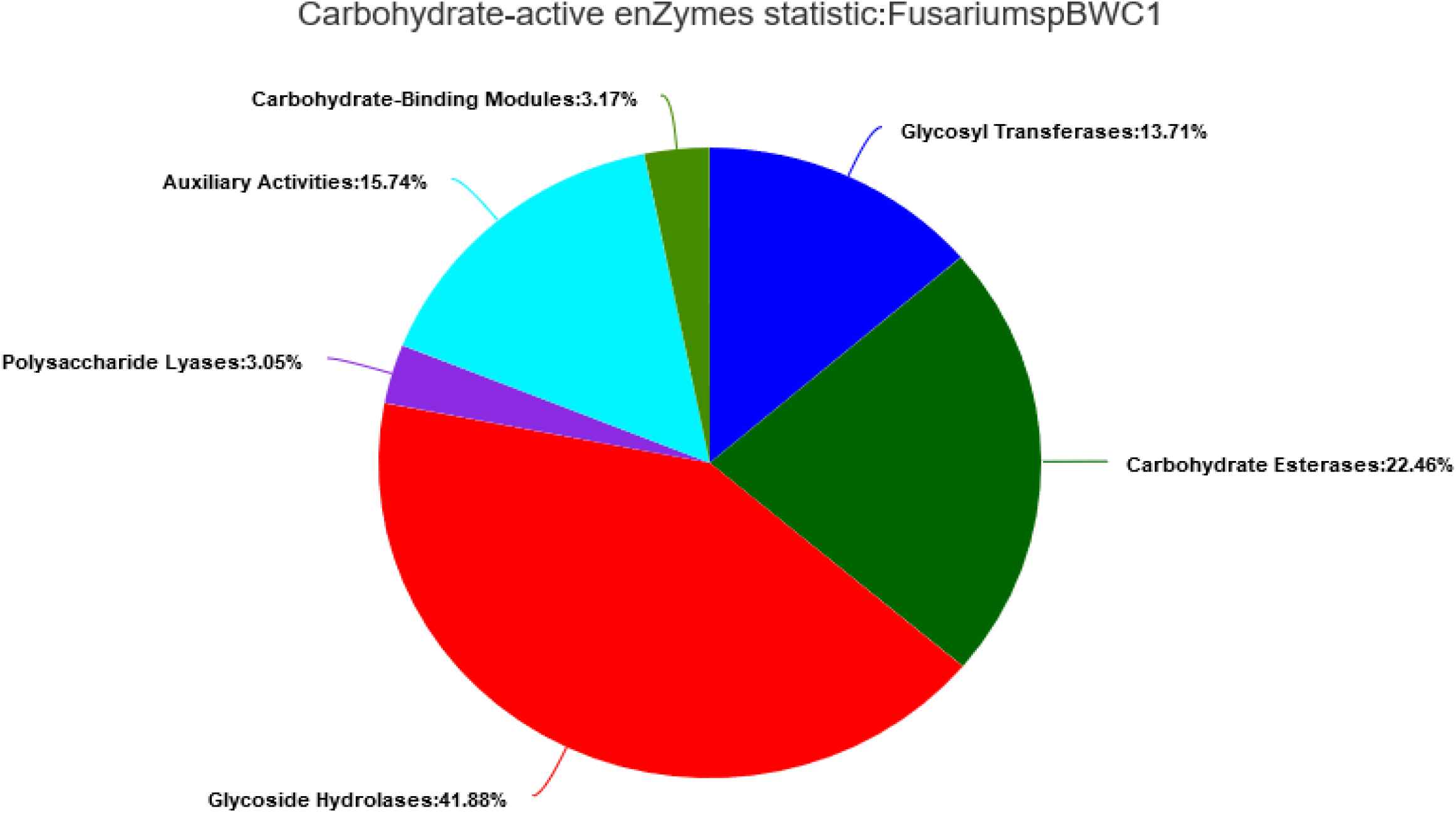
*Fusarium* sp. BWC1 CAZy analysis chart.

### 5. Phylogenetic analysis

Simultaneous use of *TUB2*, *TEF* and *RPB2* genes effectively overcame the conservativeness of *ITS* in the phylogenetic analysis. Based on the evolutionary tree shown in Figure 6, it was predicted that *Fusarium* spp. QHM and BWC1 are essentially the different *Fusarium* species that were classified into distinct clades. *Fusarium* sp BWC1 and *Fusarium subglutinans* NRRL 2016, a type strain, coexisted in same one clade and were in the closest phylogenetic relationship other than any species. Thereof *Fusarium* sp BWC1 was predicted to be a strain of *Fusarium subglutinans* at bootstrap value 1.00*. Fusarium pininemorale* CBS 137240, *Fusarium marasasianum* CBS 137238, *Fusarium parvisorum* CBS 137236 and *Fusarium fracticaudum* CBS 137234 exhibited close phylogenetic relationship with it in the *Fusarium fujikuroi* species complex (FFSC). *Fusarium* sp. QHM contained in a separated clade shown in bold font distinctly differ from other taxa members of *Fusarium.* Thereof *Fusarium* sp. QHM was putative new species of FFSC. *Fusarium oxysporum* CBS 77497 is considered as a non-FFSC or outgroup species. Taken together, the multiple loci in the phylogenetic tree topologies shown in Figure 6 are in concordance with previously reported findings for FFSC (4, 24).

**Figure 6.**
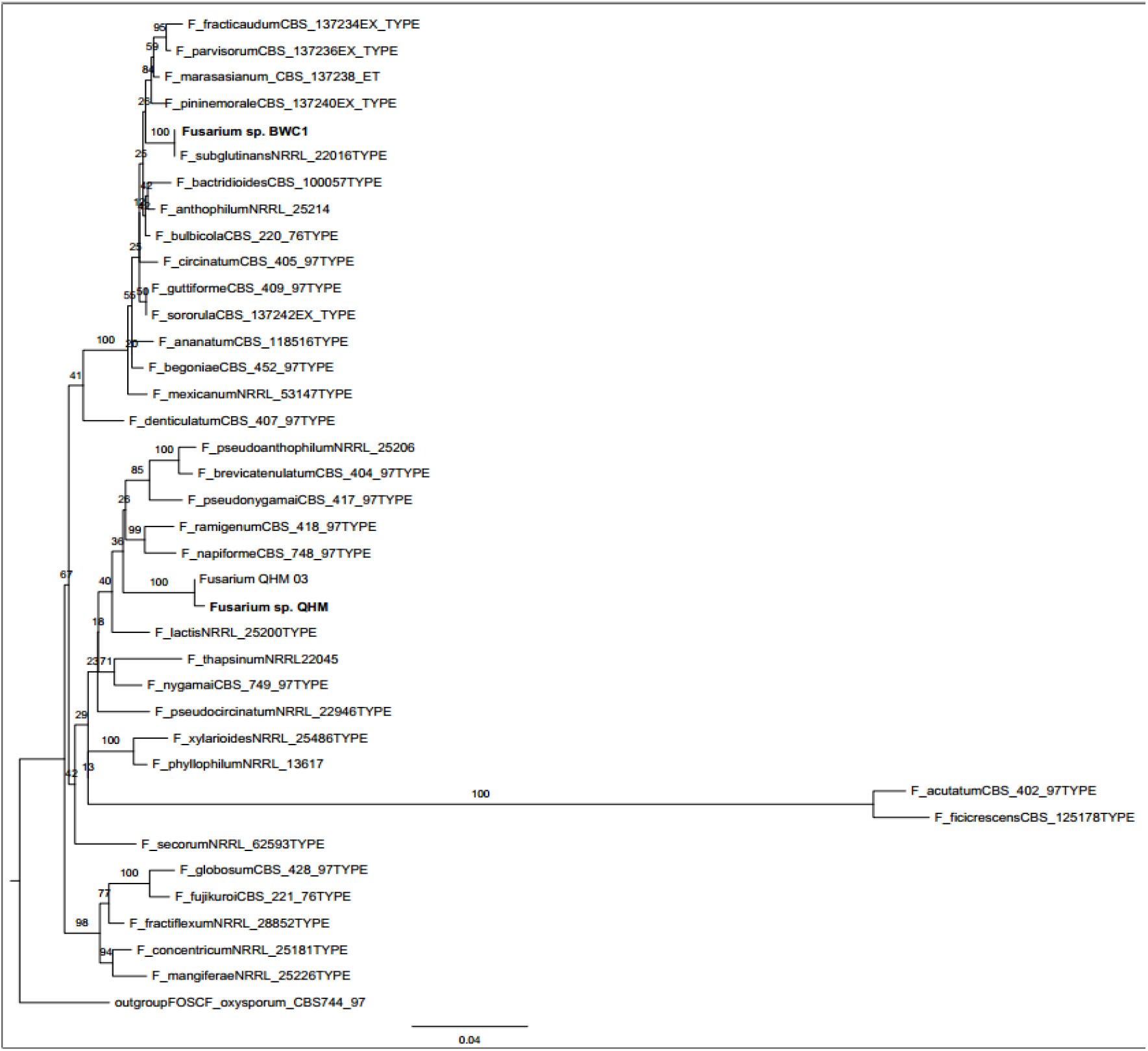
Maximum likelihood phylogeny tree. The tree was constructed based on analysis of the TUB2, TEF and RPB2 genes of *Fusarium* spp. QHM and BWC1 of the *Fusarium fujikuroi* species complex (FFSC). The names in bold font represent the two analyzed strains, that is, *Fusarium* sp. QHM and *Fusarium* sp. BWC1. Bootstrap values are provided at the branch nodes. *Fusarium oxysporum* CBS 774 was designated as an outgroup species in the analysis.

### 6. Genes associated with virulence and pathogenicity

The virulence and pathogenicity genes were identified by referencing the genes deposited in the fungal virulence factors (DFVF), comprehensive antibiotic research (CAR) and pathogen host interactions (PHI) databases. A total of 107 genes of *Fusarium* sp. QHM and 106 genes of *Fusarium* sp. BWC1 were found to play a role in the efflux pump and confer resistance against antibiotics such as glycopeptides, rifampin and tetracycline (17). The efflux pump is the main mechanism for antibiotic resistance adopted by these two fungi. Other underlying mechanisms included molecular bypass, enzymatic antibiotic inactivation, alterations in the antibiotic target, cell permeability, horizontal transfer of resistance genes and modulation of antibiotic efflux. Comprehensive data related to the antibiotic resistance mechanisms of both strains are provided in Figure 7. With regard to pathogenicity, 1487 and 1494 unaffected pathogenic genes were identified in *Fusarium* sp. QHM and *Fusarium* sp. BWC1, respectively, while 1197 and 1219 reduced virulence genes were identified. The virulent genes of clinical interest may be selected as candidate targets for control measures and novel way detection.

**Figure 7.**
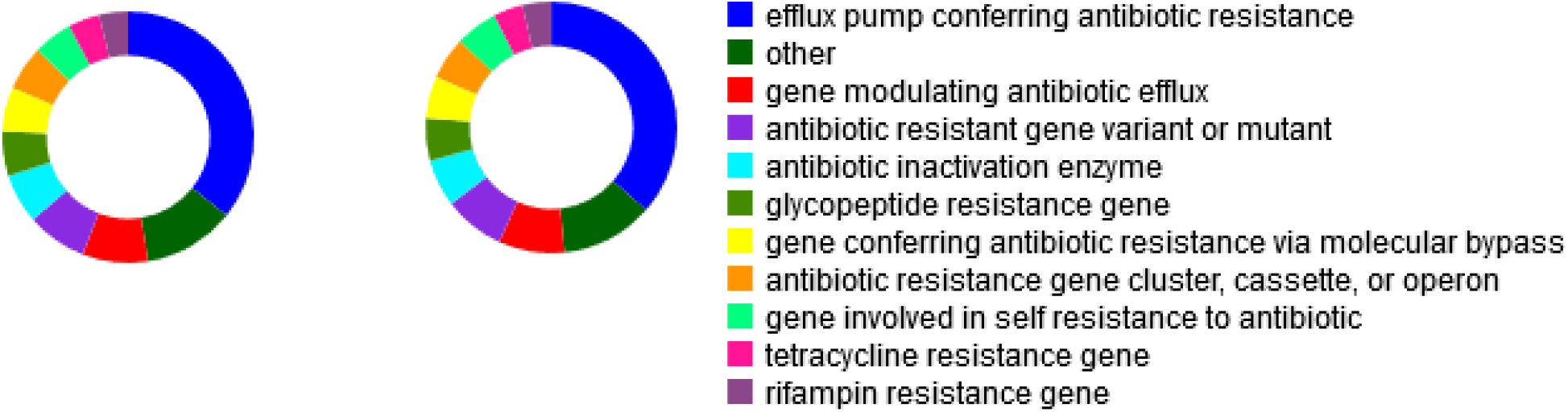
Comprehensive data on antibiotic resistance in *Fusarium* spp. QHM and BWC1. Overall mechanism of the antibiotic resistance were similar for both strains. 107 genes of *Fusarium* sp. QHM (left) and 106 genes of *Fusarium* sp. BWC1 (right) conferred the antibiotic resistance in a way of efflux pump.

### 7. Xenobiotic degradation pathway analysis

Genes involved in the degradation of a variety of xenobiotics were identified in the KEGG pathways. Based on the findings, *Fusarium* spp. QHM and BWC1 were considered to be responsible for the bio-degradation or catalysis of aromatic compounds such as chloroalkane, styrene, atrazine, chlorocyclohexane and dioxins. A detailed observation of these pathways might reveal the presence of relevant enzymes such as 2-haloacid dehalogenase, carboxymethylenebutenolidase, catechol-1,2-dioxygenase, salicylate hydroxylase, DDT-dehydrochlorinase, and biphenyl-2,3-dioxygenase. The key enzymes involved in the degradation pathway and the KEGG orthology identifiers are summarized in Table 4.

**Table 4.**
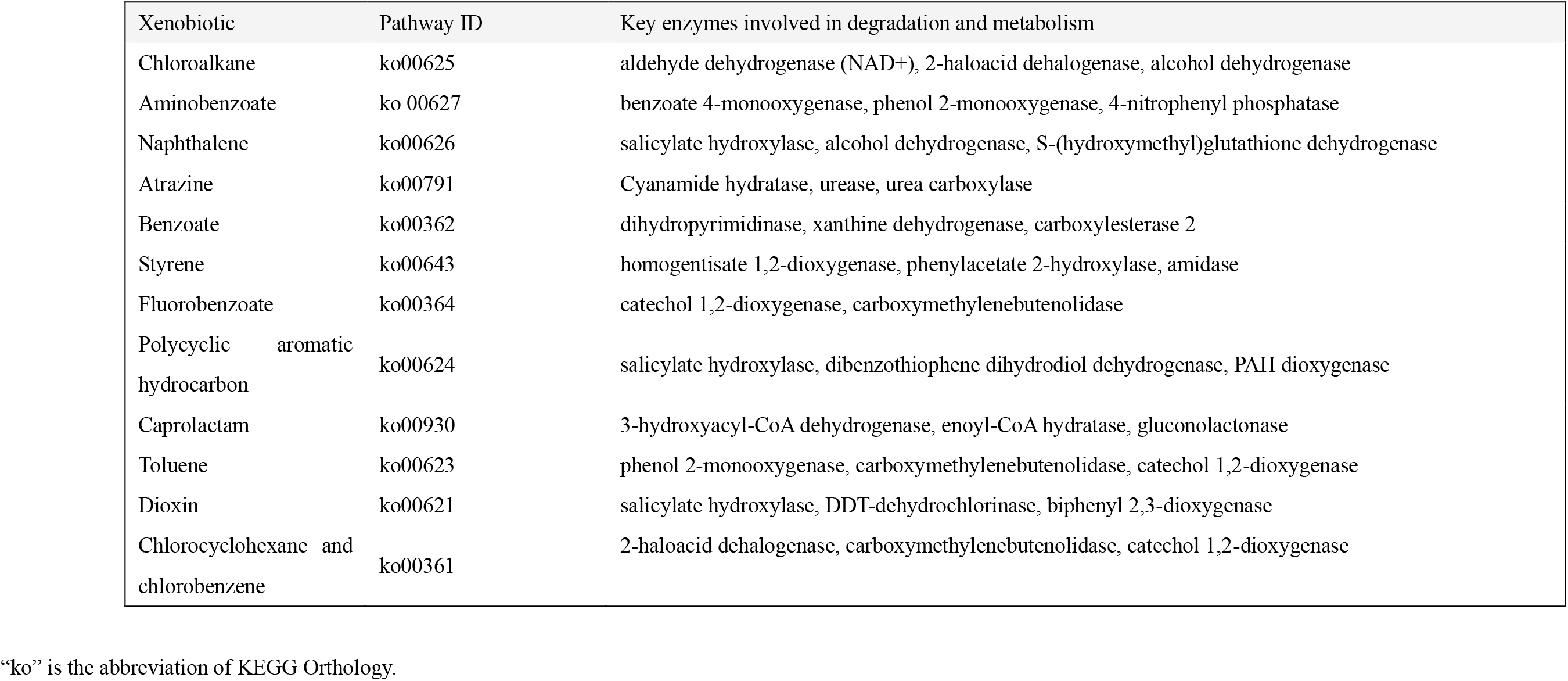
Xenobiotic biodegradation pathways and key enzymes of *Fusarium* spp. QHM and BWC1

| Xenobiotic | Pathway ID | Key enzymes involved in degradation and metabolism |
| --- | --- | --- |
| Chloroalkane | ko00625 | aldehyde dehydrogenase (NAD+), 2-haloacid dehalogenase, alcohol dehydrogenase |
| Aminobenzoate | ko 00627 | benzoate 4-monooxygenase, phenol 2-monooxygenase, 4-nitrophenyl phosphatase |
| Naphthalene | ko00626 | salicylate hydroxylase, alcohol dehydrogenase, S-(hydroxymethyl)glutathione dehydrogenase |
| Atrazine | ko00791 | Cyanamide hydratase, urease, urea carboxylase |
| Benzoate | ko00362 | dihydropyrimidinase, xanthine dehydrogenase, carboxylesterase 2 |
| Styrene | ko00643 | homogentisate 1,2-dioxygenase, phenylacetate 2-hydroxylase, amidase |
| Fluorobenzoate | ko00364 | catechol 1,2-dioxygenase, carboxymethylenebutenolidase |
| Polycyclic aromatic hydrocarbon | ko00624 | salicylate hydroxylase, dibenzothiophene dihydrodiol dehydrogenase, PAH dioxygenase |
| Caprolactam | ko00930 | 3-hydroxyacyl-CoA dehydrogenase, enoyl-CoA hydratase, gluconolactonase |
| Toluene | ko00623 | phenol 2-monooxygenase, carboxymethylenebutenolidase, catechol 1,2-dioxygenase |
| Dioxin | ko00621 | salicylate hydroxylase, DDT-dehydrochlorinase, biphenyl 2,3-dioxygenase |
| Chlorocyclohexane and chlorobenzene | ko00361 | 2-haloacid dehalogenase, carboxymethylenebutenolidase, catechol 1,2-dioxygenase |

“ko” is the abbreviation of KEGG Orthology.

**Figure 8.**
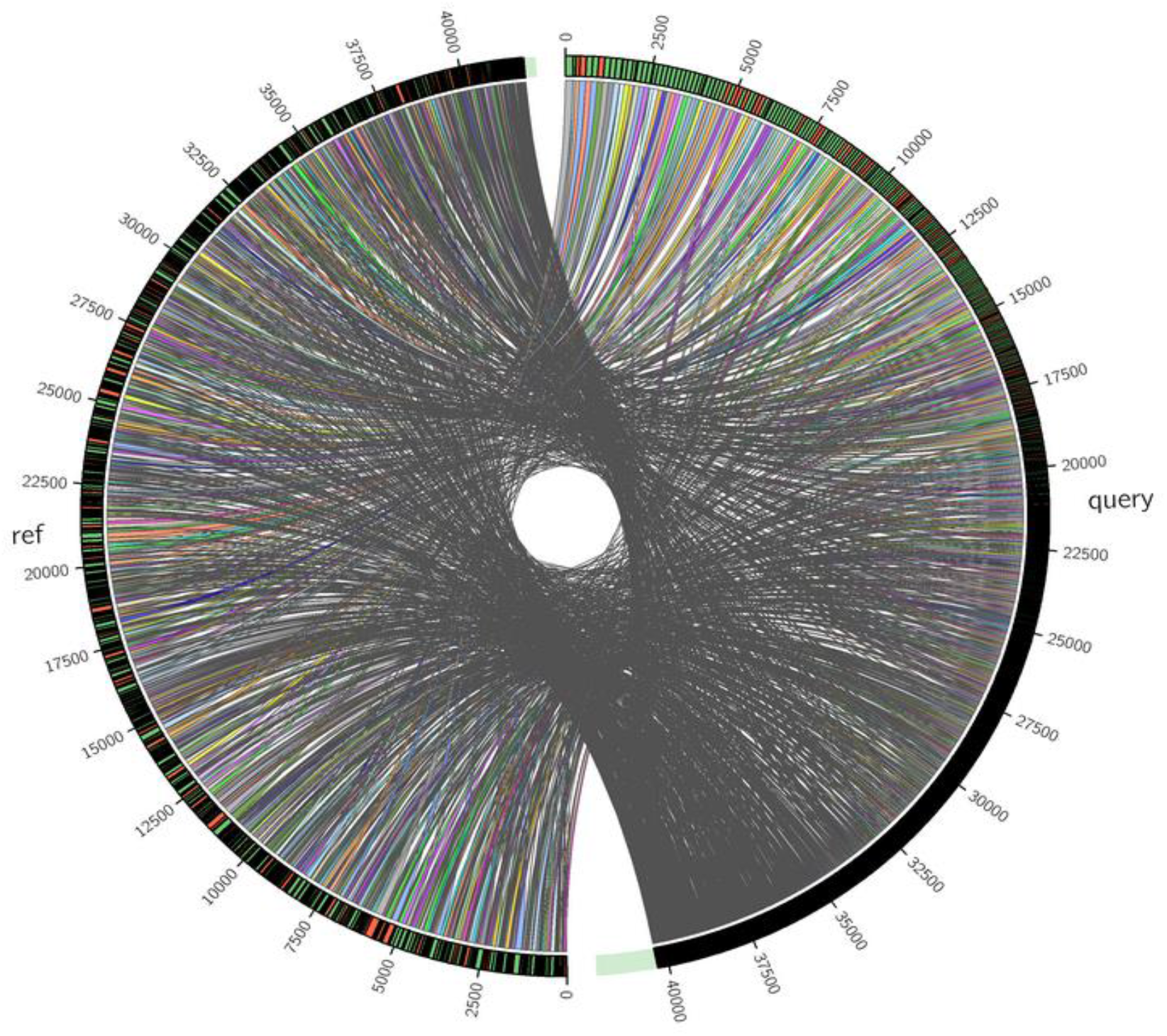
Syntenic analysis of the *Fusarium* spp. QHM and BWC1 genomes. “query” stands for QHM genome; “ref” stands for BWC1.

### 8. Syntenic analysis

The dense ligatures and concentrated genes in the DNA fragment alignments between the genomes of *Fusarium* spp. QHM (reference) and BWC1 (query) indicate a high degree of synteny, which is believed to be derived from the sequence conservation. This finding indicates the close phylogenetic relationship between the two strains, even though they were contained in distinct clade within FFSC in Figure 6. The difference in homology and orthology was proposed to be responsible for the individual adaption mechanism. Gene loss and horizontal gene transfer may somehow occurred under the natural selection pressure in the evolution.

## Discussion

The present study aims to present the draft genome sequences of the previously isolated *Fusarium* spp. QHM and BWC1 from acidic aquatic environments. The genomes were analyzed to identify the main xenobiotic degradation and pathogenic mechanisms of these species. Moreover, the phylogenetic relationship between the two species and other species in the genus was elucidated. The two species were found to be closely related phylogenetically. *Fusarium* spp. QHM and QHM 03 forms a separate clade (based on the degree of synteny) within FFSC.

Here, functional analyses of the genes and sequences revealed that the majority of the metabolism-associated genes in both species were involved in xenobiotic biodegradation, carbohydrate metabolism (the main enzyme component being carbohydrate hydrolases) and amino acid metabolism. Further analysis of the xenobiotic degradation pathways showed that *Fusarium* spp. QHM and BWC1 played a role in the bio-degradation or catalysis of several important aromatic compounds. The xenobiotic degradation ability of *Fusarium* spp. QHM and BWC1 would be potentially useful for the treatment of water that is polluted by anthropogenic activities. The use of these fungal species for tackling the pollution of water and soil would also be more effective and safer than the use of other chemical approaches. Further, the plasticity observed in *Fusarium* spp. QHM and BWC1 to adapt to low pH conditions suggests that it can be utilized in acid water control and heavy metal removal from the environment. Thus, *Fusarium* spp. QHM and BWC1 would be highly useful as bio-control agents within the indigenous microbial community in coal mine water.

There are some limitations to this study on the degradation analysis of xenobiotic, due to the lack of experimental evidence. It has been well documented that the cytochrome P450 monooxigenase plays pivotal role in the degradation process by the oxygen atom addition to the bonds between carbon and hydrogen, or among the carbon atoms. Future studies should focus on the investigation of P450 transcriptomics of these two isolates in different pH and aromatic concentration.

With regard to their pathogenic mechanisms, the present findings revealed that the efflux pump was the major mechanism of antibiotic resistance commonly adopted by both bacterium and fungus. The efflex pump components should be taken as drug targets in the fungal disease control or prevention. It has been reported that *Fusarium subglutinans* may function as a maize pathogen and the phyto-virulence is in need of further investigation of this strain. The potential infectivity and pathogenicity of the *Fusarium* community, especially for the environmental resource are to be fully aware of.

Multigene phylogenetic analysis generally provides stronger resolution than analysis of a single conservative gene. The identity of ITS shared by *Fusarium* spp. QHM and BWC1 is 100%, which makes it necessary to adopt to multigene phylogenetic analysis. Under the analysis, *Fusarium. pseudonygamai* CBS 41897, *Fusarium pseudoanthophilum* NRRL 25206, *Fusarium lactis* NRRL 25200 type strain, *Fusarium brevicatenulatum* CBS 40497, and *Fusarium napiforme* CBS 74897 were found to be closely related to *Fusarium* sp. QHM in FFSC. The two *Fusarium* species were well separated from the others and formed a separate clade (QHM) with high bootstrap support for three or four-gene datasets. Therefore, multigene phylogenetic analysis provides genealogical exclusivity for molecular fungal taxonomy. Moreover, genomic information may provide insight into the growth, spread, and infection control of this fungus. In particular, information about the genome sequence of *Fusarium* spp. QHM and BWC1 would have important implications for aquatic quarantine and water quality assessment.

In conclusion, the findings of this study have important implications for future research on the potential use of these species as biocontrol agents for the treatment of polluted water bodies and soil. The genomic information provides opptunity and tool for the control of their pathogenicity or the etiological detection.

## Materials and methods

### 1. Isolation and cultivation

Water samples were obtained from Fuxin Qinghemen (for QHM) coal mine, which is located 700 m underground, and Benxi water cave (for BWC), Liaoning, China. Both the QHM and BWC sampling sites are 20 cm below the river water surface. Each water sample (5 mL) was added to 100 mL of 9 KG liquid medium at pH 3.0–3.5, and this solution was used for *A. cryptum* cultivation. The 9 KG medium has the following composition: (NH_4_)_2_SO_4_, 3.0 g; K_2_HPO_4_, 0.5 g; MgSO_4_⋅7H_2_O, 0.5 g; Ca(NO_3_)_2_, 0.01 g; tryptone, 0.1 g; C_6_H_12_O_6_, 1 g (sterilizing filtered); Agar, 20 g (if solid) in 1000 mL (8, 9). The pH of the solid medium was 4.5. All the ingredients (with the exception of glucose) were sterilized for 20 min, as per the standard protocol.

The inoculated flask was shaken to turbidity at 100 rpm (rotation per minute) at 30°C for 24–36 h. Subsequently, the liquid culture was spread on a solid plate so as to enable the isolation of a single colony. The single colony, which appeared white, smooth and round (approximately 1 mm in diameter), was selected for enrichment on the solid medium. The solid culture derived from the single colony was inoculated in a liquid culture to increase growth for identification and storage.

### 2. Characterization

A series of 9 KG liquid media, across a pH gradient of 1.0, 2.0, 3.0, 4.0, 5.0, 6.0, 7.0 8.0 and 9.0, was prepared to investigate the acidic tolerance of the strains under the same cultural conditions as described above. Culture growth under the gradient pH was manually observed.

Polymerase chain reaction (PCR) targeting the ITS1, TUB2, TEF and RPB2 genes was performed primarily for species identification through BLAST (Basic Local Alignment Search Tool) with the reference genes obtained from the relevant public databases. The PCR primers used in the present work and their sequences are provided in Table 1.

### 3. Multigene phylogenetic analysis

Since the ITS sequence is known to be relatively well conserved within the fungal clade of the species complex, the phylogenetic relationship of the two isolates with other species in the *Fusarium* genus was analyzed based on multiple gene loci, including TUB2, TEF and RPB2 (10–12). The simultaneous use with multiple gene loci may reduce the phylogenetic bias and provide a comprehensive understanding of the phylogenetic relationships (3). The PhyML software was used to construct the maximum-likelihood tree for the TUB2, TEF and RPB2 gene sequences with the neighbor-joining method (13).

### 4. Genome sequencing

Genomic DNA extraction for *Fusarium* spp. QHM and BWC1 was carried out with the universal DNA purification kit from Tiangen Biotech (Beijing) Co. Ltd. Shot-gun sequencing on the Illumina Hiseq×10 Platform was conducted by Shangai Majorbio.

### 5. Sequence assembly and gene function prediction

SOAP denovo v.2.04 (http://soap.genomics.org.cn/) was used for de novo assembly of the short sequences obtained from the clean data. Trimmomatic, SeqPrep, Sickle and FastqTotalHighQualityBase.jar etc. had been used to acquire clean data. Maker 2, Barrnap 0.4.2, tRNAscan-SE v1.3.1 and RepeatMasker were used to predict the gene function of coding sequences, rRNA, tRNA genes and repeat sequences. All these softwares were provided by the i-sanger cloud platform (https://www.i-sanger.com/).

### 6. Gene annotation

The first step is fundamental annotation based on sequences and functions deposited in the five most well-known databanks, i.e., Non-Redundant Protein Database (NRPD), Swiss-Prot, Pfam, Clusters of Orthologous Groups of proteins (COG) and GO. The gene function of the coding sequences was predicted by means of BLAST (14, 15).

In the next step, KEGG analysis was used to identify the pathways that the genes were involved in; the encoded proteins were primarily analyzed using the Diamond software. The analysis includes pathways involved in metabolism, diseases, organismal systems, genetic information processing, environmental information processing and cellular processes.

With regard to carbohydrate metabolism, the carbohydrate active enzymes (CAZy) database with the Diamond software was used (16). For pathogenicity and antibiotic resistance, DFVF, CAR and PHI databases were used to determine the functions of virulent or resistant genes (17, 18). The Diamond software was extensively used to identify the virulence genes, pathogen–host interactions, secretion proteins, transport proteins and trans-membrane proteins through alignment of the obtained genomes (16, 19).

### 7. Bioinformatics-based analysis of the genes involved in the degradation of xenobiotics

From the environmental perspective, the metabolic pathways of aromatic compounds, such as chloroalkane and styrene, are of importance (20). A variety of enzymes present in these pathways play a role in the degradation of xenobiotics by the *Fusarium* spp. QHM and BWC1. The key enzymes involved in xenobiotic degradation were manually identified based on the reference pathways in the KEGG database (21–23).

### 8. Syntenic analysis of the genomes of the *Fusarium* spp. QHM and BWC1

Syntenic analysis, which is one of the methods of comparative genomics, is the main method used to investigate homology between species. The sequences of the *Fusarium* sp. QHM, which served as a reference, were aligned against those of the *Fusarium* sp. BWC1 to determine the homology and orthologues between the species. The orthologous and homologous regions between the contigs of the two speci were aligned to make a contiguous block within the range. The Sibelia and Circos softwares installed in i-sanger platform were used to carry out the comparison.

## Acknowledgments

This research did not receive grants from any funding agency in the public, commercial, or not-for-profit sectors.

